# Genome-wide mutational diversity in *Escherichia coli* population evolving in prolonged stationary-phase

**DOI:** 10.1101/098202

**Authors:** Savita Chib, Farhan Ali, Aswin Sai Narain Seshasayee

**Affiliations:** National Center for Biological Sciences, Bengaluru, Karnataka, India-560065

**Keywords:** Prolonged stationary-phase, Long-term stationary-phase evolution (LTSPE), experimental evolution, population diversity

## Abstract

Prolonged stationary-phase is an approximation of natural environments presenting a range of stresses. Survival in prolonged stationary-phase requires alternative metabolic pathways for survival. This study describes the repertoire of mutations accumulating in starving *E. coli* populations in lysogeny broth. A wide range of mutations accumulate over the course of one month in stationary-phase. SNPs constitute 64% of all mutations. A majority of these mutations are non-synonymous and are located at conserved loci. There is an increase in genetic diversity in the evolving populations over time. Computer simulations of evolution in stationary phase suggest that the maximum frequency obtained by mutations in our experimental populations can not be explained by neutral drift. Moreover there is frequent genetic parallelism across populations suggesting that these mutations are under positive selection. Finally functional analysis of mutations suggests that regulatory mutations are frequent targets of selection.

## Introduction

Microorganisms often face difficult conditions, including nutrient limitation. The model bacterium *Escherichia coli* doubles its population every thirty minutes during exponential phase in rich laboratory medium. This is in contrast to Savageau’s estimate that in its natural environment - predominantly the lower gut of warm blooded animals - the average doubling time of *E. coli* might be as long as 40 hours [1]. Further, in their natural environments, in contrast to standard laboratory media, bacteria are often exposed to a variety of other stresses including pH and oxidative stress [2]. In addition, many natural environments are fluctuating in their nutrient content as well as in their presentation of other stresses. Such environments constantly select for genetic variants that are better adapted to the prevailing conditions than their parents were, thus driving evolution. A particular laboratory model for approximating stressful and dynamic environments - characterised by a heterogenous population of a bacterial species - is prolonged stationary phase [3,4].

In a typical batch culture of *E. coli* maintained in a controlled environment, bacterial cells divide rapidly and quickly exhaust readily available nutrients [5,6]. Following a brief exponential growth phase, the population makes a transition to the stationary phase wherein resources are diverted to maintenance and survival rather than growth and population expansion [7]. After 48 hours in stationary phase, the medium is unable to support large population sizes, resulting in a crash in which ~99% of the cells in culture die. What survives will include mutants that are able to grow under nutrient limiting conditions. As cells die and lyse, novel nutrient forms become available, and variants which are able to metabolize these better are selected for. This is a continuous process and may extend over a period of several years. This phenomenon is termed as GASP (<u>G</u>rowth <u>A</u>dvantage in <u>S</u>tationary <u>P</u>hase), and mutations that confer growth advantages in extended stationary phase are referred to as GASP mutations. The GASP phenomenon was demonstrated in *E. coli* first and has been shown in other bacteria as well [8].

The growth advantage conferred by a GASP mutation is typically demonstrated by mixed culture competition experiments in which a mutant is directly competed against the parent under stationary-phase. General trends that have emerged from such GASP studies include the following: (a) a wide spectrum of mutations is selected in a short period of time resulting in rapid adaptation [8–11]; (b) nutrient limitation is a predominant force of selection as mutants with enhanced ability to scavenge amino acids and DNA from the debris display GASP phenotypes [10,12–14]; (c) there is increased phenotypic diversity, as reflected by colony characteristics, in the population over time [15]; (d) the ever changing biochemical composition of the population continuously redefines the niche [4]; (e) global regulators of transcription are frequent targets of mutation [8,9,14]. Despite these studies, the genetic composition of a population of *E. coli* over prolonged stationary phase and its dynamics remain incompletely catalogued and understood.

In this study, we systematically explored the population diversity emerging in E. coli populations evolving for 28 days in a lysogeny broth (LB) batch culture incubated without additional nutrient supplementation. Using whole genome sequencing of population genomic DNA, we catalogue the rise and fall of multiple mutations during prolonged stationary phase, assess the extent to which the observed genetic diversity could be explained by neutral drift, and test for parallelism in the rise of mutations across multiple evolving lines.

## Methods

### Evolution experiment

Six biological replicate cultures of *E. coli* K12, strain ZK819 - a derivative of W3110 - were seeded at a 1:1000 dilution in lysogeny broth (LB), from overnight grown cultures. The starting culture volume was 200 ml in a 500 ml capacity flask. Incubator shaker was set at 37°C and 200rpm. To compensate for evaporation, appropriate amounts of water was added at regular intervals. Population size for each replicate was tracked by serial dilution plating of 0.1 ml of the culture regularly during the entire duration of the experiment. 1 ml culture was drawn from the evolving population each day, and duplicate glycerol stocks - for storage at -80°C - were made. Periodically a sample was drawn from each evolving population to extract genomic DNA for whole genome re-sequencing.

### Sequencing & Data analysis

Genomic DNA was isolated from a sample of the evolving population at days zero, two, four, six, ten, sixteen, twenty-two and twenty-eight. The isolation was performed using the GenElute^TM^ Bacterial Genomic DNA kit (Sigma Aldrich), following the manufacturer’s protocol. Sequencing libraries were prepared using Illumina’s TruSeq Nano DNA LT library preparation kit. Whole genome sequencing was performed at Center for Cellular and Molecular platform (C-CAMP) on the Illumina HiSeq1000 platform according to manufacturer’s instructions. For each sample, an average of about 13 million 101-base reads were generated, resulting in a coverage of 300X.

The reads were mapped to the reference *E. coli* genome W3110 (GenBank accession NC_007779.1), and putative SNPs, small indels and structural variants were called using the breseq pipeline, which uses Bowtie for sequence alignment [16,17]. Variant calling for the founder population was done to identify pre-existing mutations. Breseq was run with its default parameters (using the -p option to identify mutations covered by only a subset of reads), and only those mutations predicted with high confidence (under the heading “predicted mutations”) were used for further analysis. Details of mutations identified through breseq for all samples can be accessed from the supplementary website: http://bugbears.ncbs.res.in/ZK819_ltsp_evol/. Mutations observed to be present at 100% frequency in all samples were considered as ancestral mutations and were excluded from the analysis. The following mutants were validated by Sanger sequencing using specific primers: *cpdA*Δ3(488-490), *cpdA*(L147P), *rpoS819_*2 and *imp*Δ21(**Table 3**). All subsequent data analyses on this set of mutations were performed using the statistical programming language R (v.3.3.0).

### Scoring functional conservation for non-synonymous mutations

Sites carrying nonsynonymous mutations in protein-coding genes were tested for their level of conservation across gammaproteobacteria. Amino acids were grouped into six physicochemical classes (aliphatic, aromatic, polar, positive, negative & special amino acids) as described earlier [18]. A mutation was classified as ***conservative*** if the substitute amino acid was in the same physicochemical class as the original. Only those mutations which were observed at a frequency higher than 5% in any sample were selected for this analysis.

Homolog search for these proteins was carried out using *phmmer* (HMMER 3.1b1) across 246 reference proteomes listed under *Gammaproteobacteria* in UniProtKB (ref, ref). The ortholog call was based on the bidirectional best hit method using an inclusion threshold of 10^−10^ [19]. A global alignment of each ortholog was performed with the query sequence using the Needleman-Wunsch algorithm implemented in EMBOSS, and orthologs that shared more than 50% identity were selected (ref; (EMBOSS:6.6.0.0)). A multiple sequence alignment was built using these orthologs and then used to score the conservation of each residue of the protein [20]. MUSCLE software (v 3.8.31) was used to build the multiple sequence alignment.

Shannon’s entropy[21], measuring the physicochemical diversity at each residue position across a protein’s orthologs, was calculated as follows:

*Shannon’s Entropy*,

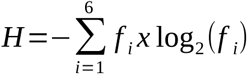

where *f_i_* represents the frequency of each of the six categories of amino acids considered here.

This was then transformed to conservation score using following expression:

*Conservation Score*,

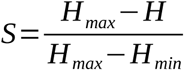

*Where H_max_ = Shannon’s entropy of the least conserved residue of the protein & H_min_ = Shannon’s entropy of the most conserved residue of the protein*

### Functional class over-representation test

A functional classification of all genes in which coding region substitutions and indels were found was done using the MultiFun classification scheme [22]. Six broad functional categories - namely metabolism, information transfer, regulation, transport, cell processes and cell structure - were considered. Only those genes listed in both MultiFun and the ‘Ecogene’ database were considered for the analysis. This excludes 81 of the 3398 genes functionally annotated in MultiFun. Out of the remaining 3317, 3041 fall in the six functional categories mentioned above and the rest were categorized as ‘Others’.

A statistical over-representation test was performed [23]. The sum total of the length of all genes in a functional category was used to get the probability of a mutation occurring in any gene of that functional category in the genome. The p-values were calculated under the binomial probability distribution for the proportion of mutations observed in genes for each functional category. The p-values were adjusted for multiple testing using the Holm-Bonferroni method.

### Genetic Diversity and models of neutral evolution

The genetic diversity was measured in terms of Shannon’s entropy[21] of the genome. Shannon’s entropy at each substitution site of the genome was calculated as:

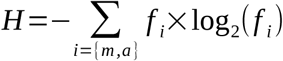

*where f_i_* represents the fraction of genomes in the population with a mutant (*m*) or the ancestral (*a*) base at the site of substitution.

The Shannon’s entropy of the genome was then calculated as:

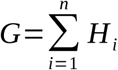

where *H_i_* represents the Shannon’s entropy at each substitution site.

And *n* represents the total number of observed substitution sites in the population.

Neutral evolution simulations were performed under two models of genetic drift in a bacterial population of constant size with no recombination or de-novo generation of novel mutations. The two models - namely the Wright-Fisher [24,25] and the Fission model [26] - differ from one another in the rate at which the frequency of one type of individuals changes over time in the population. For both models, the change in genetic diversity over time was simulated under a two-type or a multi-type system, as defined below.

In a two-type system, all observed mutations in the population constitute a single haplotype, such that an individual is either the mutant or the ancestor. The change in frequency of the mutant over a span of 20 days, if initially present at 1%, was simulated based on the rate of change in frequency governed by the model. The simulation was repeated 1000 times. The fraction of simulations in which the mutant attained a frequency of 5% or above was defined as being equivalent to the p-value for the observed genetic diversity.

In a multi-type system, mutations are distributed over several haplotypes. If the number of substitutions is much smaller than the population size, then each substitution is likely to represent a unique haplotype. The overall change in entropy was simulated by building an ensemble of genomes of length equal to the number of substituted sites. The simulation was begun by randomly selecting 1% of genomes to carry a substitution at each site. The ensemble was rebuilt after every generation based on the number of copies each haplotype would leave for the next generation. Any site with a mutant frequency lower than 1% was considered to have zero entropy. To compensate for the loss of entropy from about half of the substituted sites, the p-value was determined by the fraction of simulations in which the ensemble attained a diversity equal to or more than half of the observed genetic diversity.

Under the Wright-Fisher Model, the number of descendants each individual leaves in the next generation is approximately Poisson distributed with an average of 1 descendant / generation. With only two types of individuals, carrying either the ancestral or the mutant allele, the frequency of occurrence of each type in any given generation can be calculated from a binomial sampling of alleles from the previous generation. The maximum number of offsprings left by any individual at the end of a generation is solely limited by the population size. Note that a generation here is not equivalent to the bacterial doubling time but to any span of time at the end of which the population is observed to be constant.

Under the Fission Model, the population size is maintained as in any generation the number of bacteria dividing is equal to the number of bacteria perishing while the rest of the population is allowed to continue to the next generation without dividing. The birth-death probability and the number of generations govern the extent of diversity a population attains in the given duration of time.

## Results

### Evolution in batch culture under prolonged stationary-phase

We allowed six replicate populations of *E. coli* K12 ZK819 to stay in stationary phase for twenty eight days without further supplementation of fresh nutrients. Using the dilution-plating method, we found that the population size of the culture did not change much, after a decline of approximately 2-logs at the 3-day mark (Fig1A). Genomic DNA was isolated from a periodically sampled population for next-generation sequencing. The conclusions drawn in this report are from four replicates. Replicate5 was eliminated because of contamination detected during third week of the evolution experiment. Replicate#4 has a mutator like phenotype (Fig 1B (◊)) and is excluded from most of the analysis, except where indicated.

**Fig 1:**
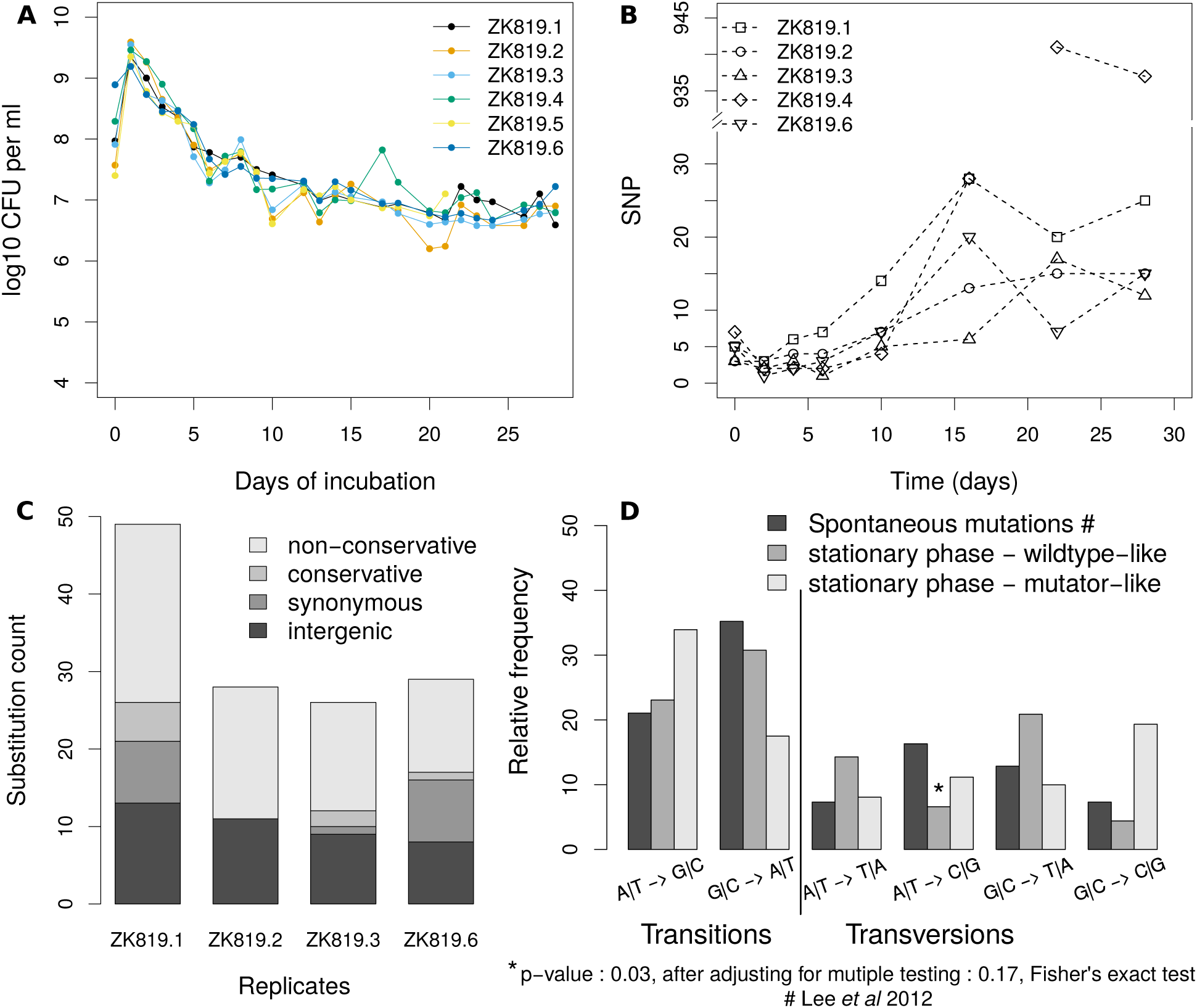
Mutation spectrum analysis in prolonged stationary-phase populations in comparison with Mutation accumulation lines (A) Change in population size over time for 6 replicate populations. (B) Change in observed number of SNPs over time for 5 replicate populations. ( ZK819.5 was discarded because of contamination detected during third week). (C) Substitution profiles of four replicates. (D) Comparison of BPS spectra from prolonged stationary phase with that reported for spontaneous mutations in a neutrally evolving population.

### Status of *rpoS819* during 28 days of evolution

ZK819 is a W3110 variant originally isolated from an *E. coli* culture evolved in Roberto Kolter’s laboratory under an environment similar to that used in our study[8]. ZK819 carries an allelic variant of the stationary-phase sigma factor gene *rpoS*, named *rpoS819*. The *rpoS819* mutation alone is sufficient to confer growth advantage to its bearer under prolonged starvation [8,12].

*rpoS819* has a 46 bp duplication at the C-terminal end, resulting in a variant that codes for a longer sigma factor with attenuated activity [8]. Genome sequencing data of the population on the first day showed heterogeneity at the *rpoS* locus. The *rpoS819* allele was found in only 70% of the population - as indicated by the proportion of reads supporting the mutation - after twelve hours of growth (Fig2). Additionally, we noted that there is a A to G transversion 21 bases upstream, and presumably in the UTR, of the *rpoS* ORF; this mutation is present at 100% frequency throughout the experiment.

**Fig 2:**
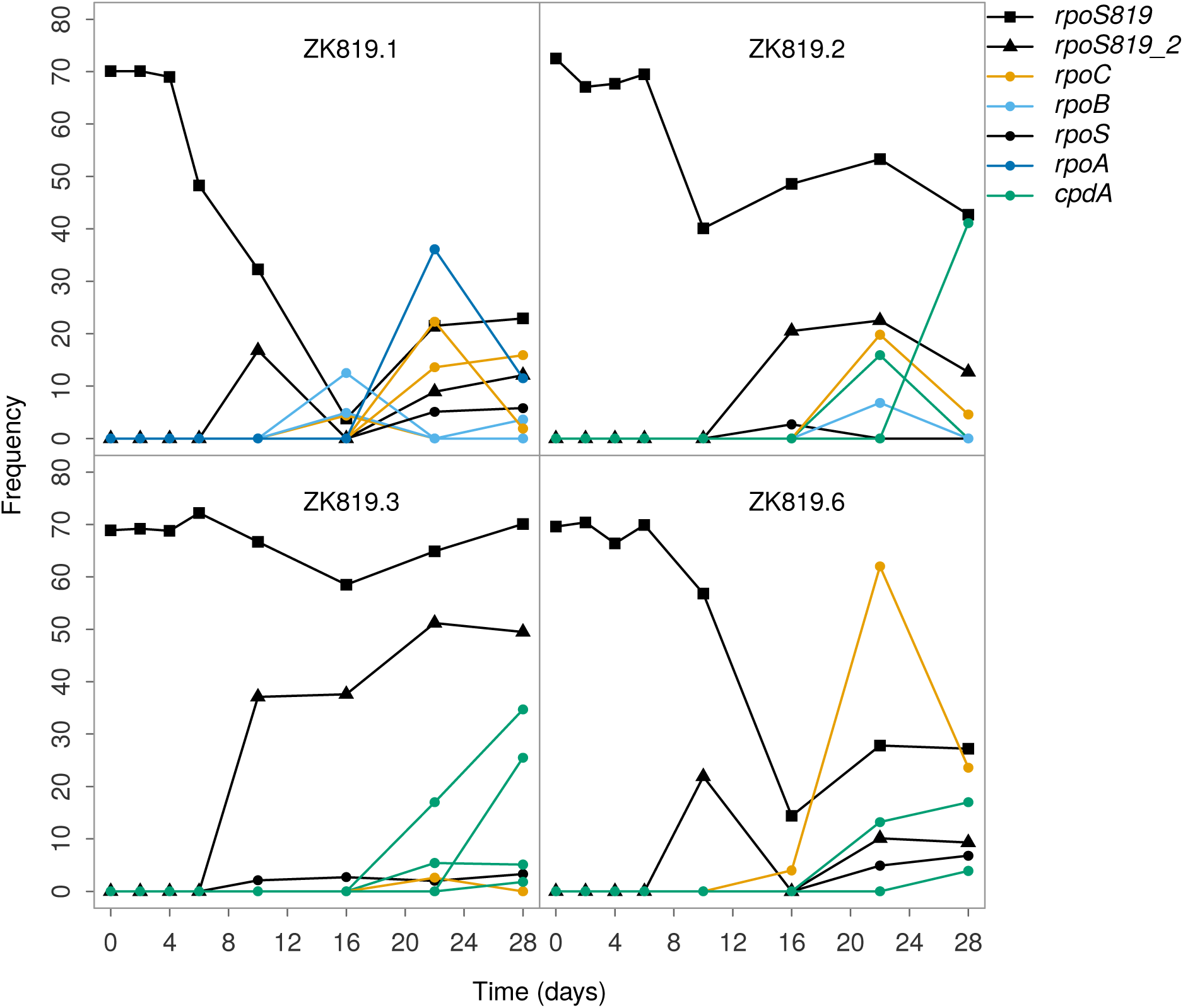
Frequency of ancestral *rpoS819*(stationary sigma factor)and new rpoS819 allele variant (*rpoS819_2*) fluctuates and exhibit different patterns across replicates(■,▲). Other genes such as *cpdA*- a cyclic AMP diphosphoesterase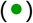 and *rpoC, rpoB* and *rpoA*- RNA polymerase subunit 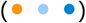 repeatedly appeared in multiple replicates.

**Fig 3:**
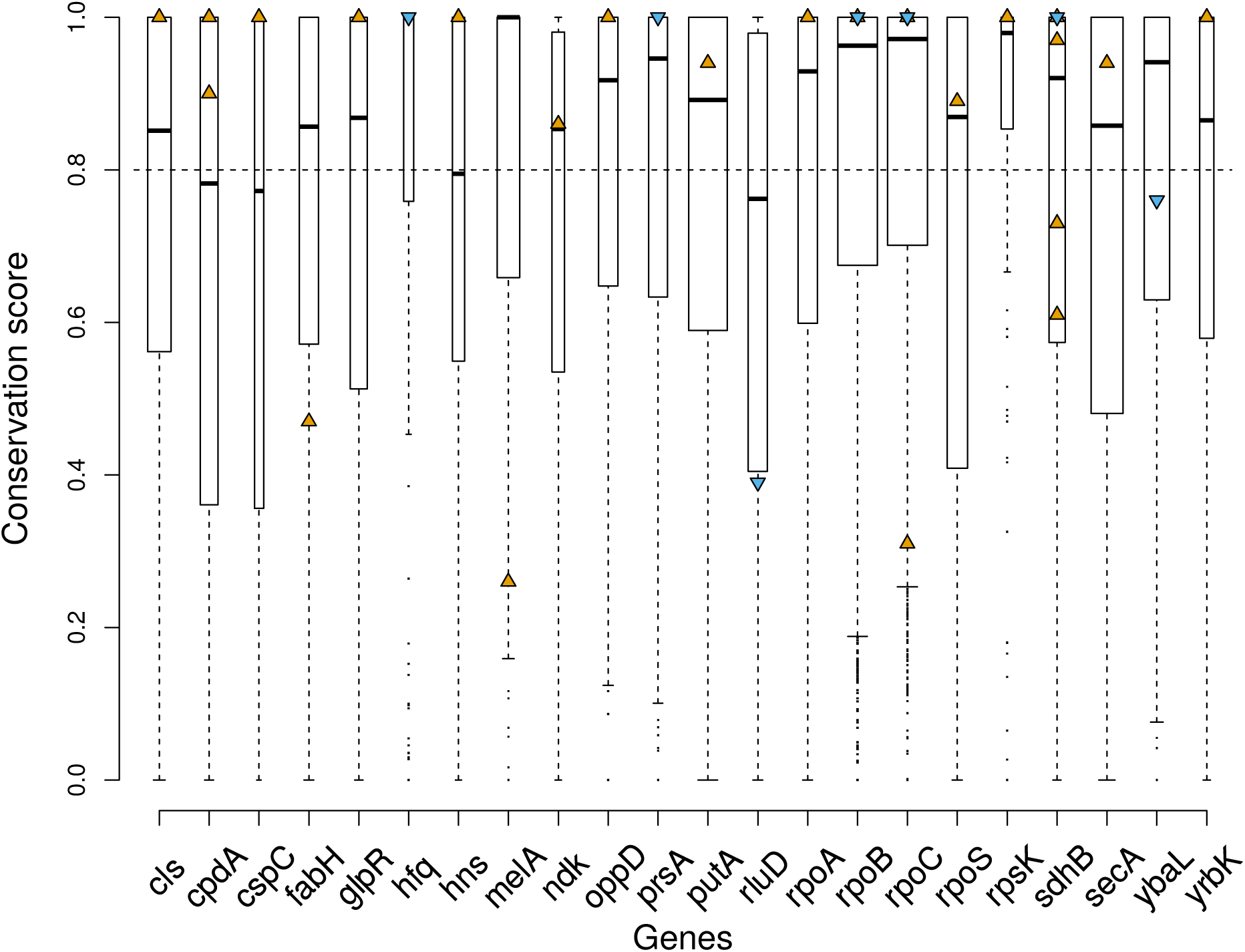
Highly conserved residues mutated in prolonged stationary phase. Conservation score was calculated for each protein using Shannon’s entropy (see methods). The width of boxes is indicative of protein sizes. Triangles mark the score for mutated residues in each protein, orange indicates a non-conservative amino acid change while blue indicates a conservative one.

**Fig 4:**
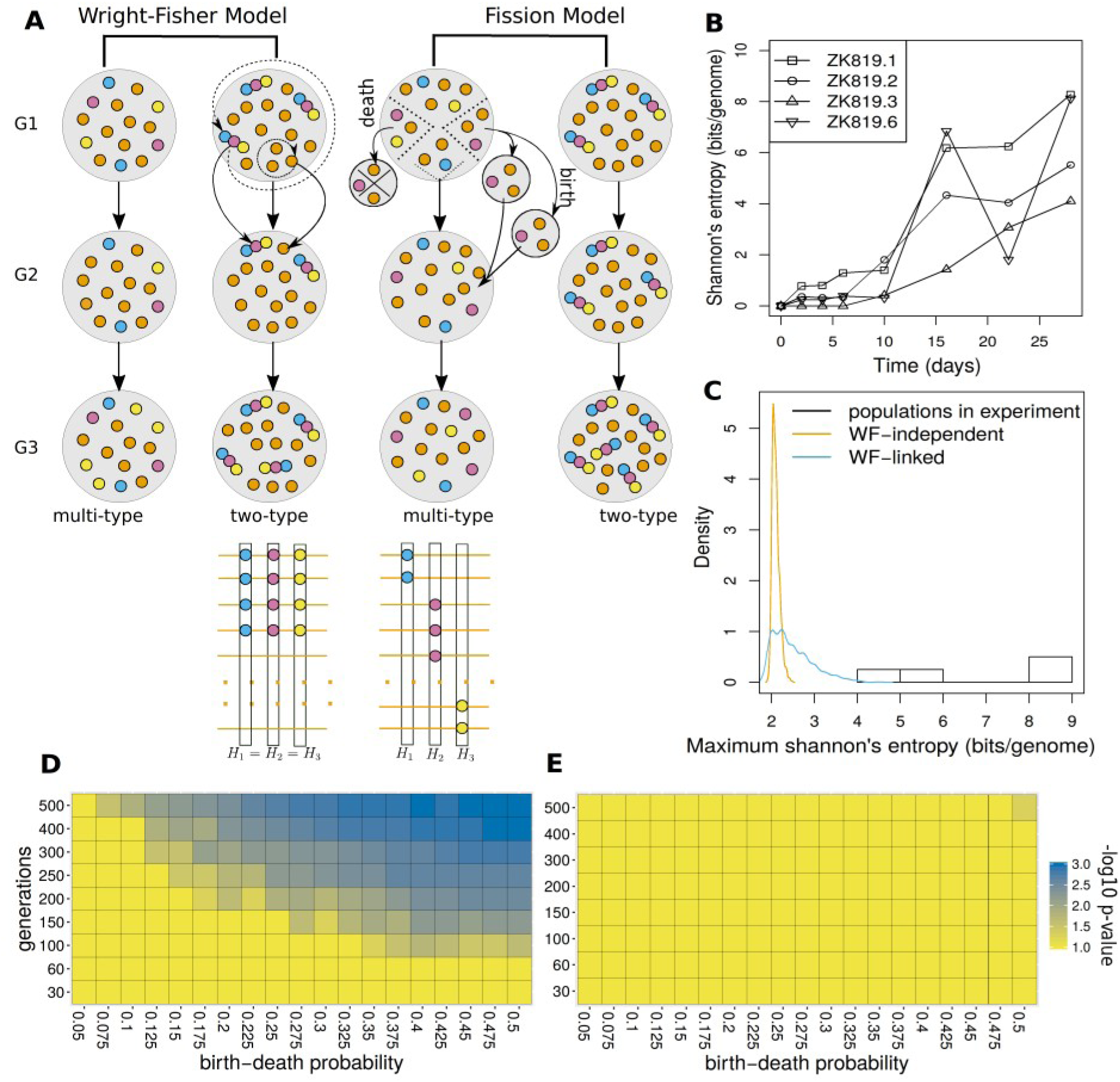
Simulation of neutral evolution in prolonged stationary phase. (A) Schematic for Wright-Fisher and Fission model used for neutral evolution simulation, illustrated by running a simulation with a population of 15 individuals for 3 generations and calculating Shannon’s entropy at each mutated position in the genome. Orange circles indicate ancestral allele, other colors indicate mutant alleles, circles stuck together indicate all mutant alleles to be present on the same genome. (B) Change in Shannon’s entropy of the whole genome (calculated from frequency of observed substitutions in the population) over time for 4 replicates. (C) Distribution of maximum Shannon’s entropy over 20 days for 1000 simulations of Wright-Fisher model in comparison with the values observed in 4 replicates. (D and E). Parameter scan for birth-death probability and number of generations in 20 days under Fission Model: (D) Two-type (E) Multi-type.

A second allele of *rpoS* was identified in all replicates from day 10 onwards. Sanger sequencing confirmed that the original 46 base-pair duplication motif has undergone another round of duplication thereby increasing the size of the duplication to 92 base-pairs. Over the course of the experiment, we found this mutant allele - designated *rpoS819_2* - in up to 30% of the population (Fig2). Thus, we suggest that the function of RpoS is tunable and that there is a strong selection favoring it under prolonged stationary-phase.

### Spectrum of mutations accumulated over 28 days

On an average, twenty-six mutations were observed in each replicate in the twenty eighth day sequencing time point, also the end of the experiment. Sixty four percent of these mutations were substitutions. Small indels and large deletions/insertions created by insertion elements constituted the remaining set of mutations (**Table 1**). In the day twenty-two sample of the ZK819.4 population, we observed a nearly 40-fold increase in substitution mutations. We therefore refer to ZK819.4 as mutator-like.

**Table. 1.** Mutation spectrum of all replicates

|  | ZK819.1 | ZK819.2 | ZK819.3 | ZK819.4 | ZK819.6 |
| --- | --- | --- | --- | --- | --- |
| Total | 66 | 44 | 39 | 1133 | 43 |
| intergenic | 22 | 14 | 13 | 29 | 13 |
| coding | 44 | 30 | 26 | 1104 | 30 |
| snp | 49 | 28 | 26 | 1102 | 29 |
| insertion | 2 | 2 | 2 | 8 | 3 |
| deletion | 7 | 11 | 7 | 19 | 7 |
| large deletion | 8 | 3 | 4 | 4 | 4 |

Spontaneous mutations in the absence of selection are expected to leave a characteristic spectrum of Base Pair Substitutions (BPS), deviations from which may indicate the presence of mutagens or the absence of specific mismatch-repair pathways or even the presence of selection. A mutation accumulation (MA) protocol is most appropriate for scanning the unbiased pool of spontaneous mutations, as it minimizes the influence of selection by its repetitive single-individual bottleneck strategy. We compared our data on BPS to that from a previously published MA study on wild-type *E.coli* [27]. The data from four replicates excluding ZK819.4 were pooled for this comparison (**Table 2**).

**Table. 2.** Base-pair substitution spectra of LTSP populations in comparison with data on spontaneous mutations in the wild-type from MA study (Lee *et al* 2012).

|  | MA lines | ZK819.1, 2, 3 & 6 | ZK819.4 |
| --- | --- | --- | --- |
| Substitutions | 233 | 91 | 1102 |
| Transitions |  |  |  |
| A\|T -> G\|C | 49 | 21 | 374 |
| G\|C -> A\|T | 82 | 28 | 193 |
| Transversions |  |  |  |
| A\|T -> T\|A | 17 | 13 | 89 |
| A\|T -> C\|G | 38 | 6 | 123 |
| G\|C -> T\|A | 30 | 19 | 110 |
| G\|C -> C\|G | 17 | 4 | 213 |

**Table. 3.** Mutations observed in regulatory genes (as classified in MultiFun)

| gene | mutation | annotation | ZK819.1 | ZK819.2 | ZK819.3 | ZK819.6 |
| --- | --- | --- | --- | --- | --- | --- |
| *cpdA* | Δ3 bp | coding (488-490/828 nt) | 0 | 1 | 1 | 1 |
| *cpdA* | G → A | S222L (TCG → TTG) | 0 | 1 | 0 | 0 |
| *cpdA* | Δ3 bp | coding (116-118/828 nt) | 0 | 0 | 1 | 0 |
| *cpdA* | T → C | L147P (CTG → CCG) | 0 | 0 | 1 | 0 |
| *cpdA* | Δ1 bp | coding (356/828 nt) | 0 | 0 | 1 | 0 |
| *cpdA* | CTC | coding (622/828 nt) | 0 | 0 | 0 | 1 |
| *csiE* | T → C | L87S (TTA → TCA) | 0 | 0 | 0 | 1 |
| *cspC* | C → T | G38D (GGT → GAT) | 0 | 0 | 1 | 0 |
| *dcuS* | G → T | L415M<br>(CTG → ATG) | 0 | 0 | 1 | 0 |
| *dgsA* | Δ1 bp | coding (680/1221<br>nt) | 0 | 1 | 0 | 0 |
| *glpR* | C → T | R38C (CGC → TGC) | 1 | 0 | 0 | 0 |
| *glpR* | C → A | S209* (TCG → TAG) | 0 | 1 | 0 | 0 |
| *hdfR* | G → A | W187*<br>(TGG → TGA) | 0 | 0 | 0 | 1 |
| *hfq* | T → A | D40E (GAT → GAA) | 0 | 1 | 0 | 0 |
| *hns* | T → A | K32N (AAA → AAT) | 0 | 1 | 0 | 0 |
| *putA* | G → T | R339S<br>(CGC → AGC) | 0 | 0 | 0 | 1 |
| *putA* | G → T | A270D<br>(GCC → GAC) | 0 | 0 | 0 | 1 |
| *rpoS819* | 46 bp | coding (817/876 nt) | 1 | 1 | 1 | 1 |
| *rpoS819_2* | 92 bp | coding (813/876 nt) | 1 | 1 | 1 | 1 |
| *rpoS* | A → G | L288P<br>(CTG → CCG) | 1 | 1 | 1 | 1 |
| *topA* | G → T | W860C<br>(TGG → TGT) | 0 | 0 | 1 | 0 |
| *ubiB* | C → A | L464L (CTG → CTT) | 0 | 0 | 1 | 0 |

Spontaneous mutations are known to exhibit a G|C -> A|T bias. The fraction of G|C -> A|T substitutions among transitions was higher than A|T -> G|C in all four populations except the mutator like ZK819.4 (Fig. 1D). The occurrence of these substitutions did not significantly differ from those reported from the MA experiments for the wild-type, except for an under-representation of A|T -> C|G (Fisher’s exact test, p-value: 0.029, significance threshold: 0.05); this significance disappears upon correcting for multiple comparisons. On the other hand, the fractions of both types of transitions and that of G|C -> C|G transversions in the mutator line ZK819.4 was significantly different from those in the wild type MA line (Fig. 1D).

The MA study had reported a shift in BPS from G|C -> A|T to A|T -> G|C in a *mutL* strain, which lacks a component of the mismatch repair pathway. We observed the same shift in ZK819.4; in addition, we also observed a dramatic increase in G|C -> C|G substitutions which was not observed in the MA experiment for *mutL*(Fig. 1D). We did not find any mutation in the Methyl-directed Mismatch Repair system in ZK819.4; however we found that the ZK819.4 population carried a non-synonymous mutation in *uvrB* (F497Y) at a frequency of ~7% on day 22.

Overall, the BPS spectrum in prolonged stationary phase populations excluding ZK819.4 was only marginally different from the BPS spectrum resulting from spontaneous mutations in wild type *E. coli* (Pearson’s Chi-squared Contingency Table Test, p-value = 0.01 - 0.05, 2000 monte carlo simulations). The BPS spectrum of ZK819.4, however, was significantly different from other populations as well as from those of the MA lines from the wild type and the *mutL E. coli* (Pearson’s Chi-squared Contingency Table Test, p-value < 2 * 10^-3^, 2000 monte carlo simulations). In the MA study, the fraction of non-synonymous substitutions was 0.69 and 0.66 respectively in the wild-type and *mutL* lines, which is close to the *apriori* random expectation. In ZK819.4, at the last two time points where an excessive number of substitutions was observed, the fraction of non-synonymous substitutions was around 0.36. For the corresponding time points, this fraction across the other replicates was 0.88 and 0.66 respectively. The variability in these numbers across time-points do not allow us to develop a firm interpretation of these results.

### Functional Conservation at Mutated Residues across Orthologs

Functional constraints on an amino acid residue in a protein are expected to keep the residue conserved across its evolutionarily close orthologs. The residue itself may however be allowed to be replaced as long as the physicochemical properties at that residue position in the protein are not altered.

Therefore, conservation of a certain residue type at any given position across a majority of a protein’s orthologs would indicate that a mutation that alters the properties of that residue might affect the activity of the protein.

With this reasoning, we scored the level of functional conservation, across gamma-proteobacterial genomes, at residue positions affected by non-synonymous mutations. We restricted this analysis to mutations appearing above 5 % frequency in the population. We searched for orthologs of proteins carrying these non-synonymous mutations across 246 gamma-proteobacteria. We defined a conservation score, ranging from 0 to 1, for each residue (see Methods). We calculated conservation scores for all the residues of a protein by normalizing their Shannon’s entropy values between 0 and 1 such that a score of 1 would represent the most conserved residue in the protein and vice-versa. For the 22 proteins considered here, we observed that the median score was around 0.8. The scores for 24 out of 31 residues observed to be mutated in our study were among the top 50 % of scores, all above 0.8. In fact, 16 of these residues had a conservation score of 1. This observation suggests that most of the mutated residues occupy well conserved positions in respective proteins.

We noted that 24 of these 31 non-synonymous mutations altered the physicochemical class of the residue concerned. Since a majority of these mutations were at conserved positions, these can be expected to be of functional consequence for their respective proteins.

### Genetic diversity in prolonged stationary phase populations

We measured genetic diversity of the population at each time point of our experiment by measuring Shannon’s entropy of the genome [21]. In each replicate we observed an overall increase in genetic diversity with time, but the rate of change at each time point varied across replicates, with some time points even showing a brief decline in the measured genetic diversity.

We tested through simulations the null hypothesis that the observed level of genetic diversity is attainable under neutral evolution with the same number of substitutions and initial mutant frequencies as for a prolonged stationary phase population. We simulated neutral evolution assuming a constant population size, which was appropriate for our prolonged stationary phase population day 10 onwards. In these simulations, new mutations did not occur *de-novo*, and all mutations were considered to have been present at a frequency of 1% at the initial timepoint. We considered ZK819#1 as the representative case for this analysis, because it has the highest number of substitutions at the last time point and the entropy at the starting time point can be explained by considering all its 25 substitutions at a frequency no greater than 1 %. The genetic diversity of ZK819#1 reaches its maximum of 8 bits / genome by the last time point, corresponding to an average mutant frequency of 5% per site.We note that our measure of genetic diversity represents an upper estimate of true diversity in the genome because all substitutions are considered completely independent; i.e., the knowledge of a substitution at one position does not provide any information about the presence or absence or the nature of a substitution at another position. The value of this measure would remain the same even in the extreme condition where all substitutions are linked. Therefore, in our neutral evolution simulations, we considered both the extreme cases: one where all the 25 substitutions are linked and another where they are independent. For the former case, we only needed to simulate change in frequency of a single allele initially present in 1 % of the population. For the latter case, we needed to observe the change in entropy in a genome of 25 independent substitutions, where initially all the substitutions were at 1 % frequency in the population. The probability of attaining the observed genetic diversity under complete linkage is obviously expected to be much higher than with complete independence. We note here that the chance that the complete linkage scenario would be applicable to our system is low, in light of the fact that the frequency profiles of many pairs of mutations are uncorrelated.

We considered a relatively smaller population of bacteria for simulations (10^4^, 1% of our population size estimates from colony counting experiments). Frequency changes become less prominent in a larger population and so, the probability of mutations attaining a high frequency in a larger population under neutral evolution would be even lower than the simulated p-values for a population of 10^4^ bacteria.

Under the Wright-Fisher Model for neutral evolution, both cases failed to attain the observed genetic diversity (p-value < 10^−3^; 1000 iterations). Even with a 4-fold increase in the number of WF generations, the p-value under complete linkage was 0.003. We then turned to the Fission model which is considered a more realistic drift model for bacterial populations. Since the number of generations covered by our evolution experiment and birth-death rate probability in prolonged stationary phase populations are unknown, we performed simulations with varying values of these two parameters. We observed an increase in the probability of finding the observed genetic diversity with increasing number of generations and birth-death probability. When all substitutions were considered independent, the p-value for the observed genetic diversity with a birth-death probability of 0.5 and 500 generations was 2 × 10^−3^; 1000 iterations. For all other parameter combinations, it fell below 10^−3^. Under the extreme, unlikely case where all substitutions are linked, the probability of observing a genetic diversity as high as in the prolonged stationary phase populations did not remain significantly low for more than 150 generations for higher values of birth-death probability.

To get an upper estimate of number of generations in prolonged stationary phase, we considered the case of a mutation that appeared on day 16 at 100% frequency while it was not observed on day 10. This mutation could either be present on day 10 but below the level of detection or could have appeared at anytime between day 10 and day 16. If we consider this mutation to have originated as late as day 13, and increased in numbers from 1 to 10^7^ in 3 days, our estimate of number of generations under logarithmic growth assumptions would be, ⅓ × log_2_10^7^ = 8 generations/day, which sums to no more than 160 generations in 20 days of prolonged stationary phase. Given this, the probability that neutral drift can explain the observed level of genetic diversity is low. Therefore, we can infer that the genetic diversity of prolonged stationary phase populations is a signature of evolution under natural selection.

### Functional Bias in frequencies of mutations observed in prolonged stationary phase

Since the mutations were randomly sampled, we asked if we observe any bias in relative proportions of functional classes represented in the pool of mutations. If we observe a bias towards a functional category, it would mean that among the observed mutations, a larger than expected proportion is sustained in that category in prolonged stationary phase.

Using the ‘MultiFun’ classification scheme, we classified all coding region mutations (N = 93, of which 10 were synonymous) into 6 functional categories based on the functions of their carrier genes. To have sufficient power in the statistical test, we pooled the mutations from the 4 non-mutator replicate populations and set the experiment-wide significance threshold to 0.01.

We found that the category ‘Regulation’ was over-represented among the observed mutations with a p-value of 0.007 whereas ‘Structure’ was observed to be significantly under-represented. Only one out of the 22 mutations in genes categorised under “Regulation” were synonymous. All mutations observed in regulatory genes have been listed in **Table 3**.

To test if spontaneous mutations have any inherent bias toward any functional category, we performed the same analysis on data on spontaneous mutations in wild type *E. coli* from a Mutation Accumulation study [27]. We observed that the over-representation of mutations in class ‘Regulation’ was only marginally significant and and there were no significant changes for other functional classes. This comparison led us to the inference that the regulatory mutations account for a larger than expected proportion of surviving mutations in prolonged stationary phase.

Five of the fifty genes observed to be mutated in prolonged stationary phase populations lacked ‘MultiFun’ annotation viz. *glcF, hdfR, csiE, bax and ychS*. We could include the first four of these in the analysis as the required functional information was available on other databases and literature. A *ychS* allele variant with 3 point mutations, of which 2 are non-synonymous, was identified in ZK819#1 at day 16 time-point at 100% frequency. *ychS* encodes for a 91 amino acid long protein for which no additional information is available in *E. coli* databases.

### Genetic parallelism during prolonged stationary phase

Among the mutations identified are a set of genes that appeared in multiple replicates during roughly the same period of the evolution experiment (Fig.5B). We have mentioned the case of *rpoS819*_2 earlier (Fig. 2, 5B). We observed that the components of the RNA polymerase are frequently targeted under prolonged starvation. *rpoC* alleles appeared in four evolving populations. Replicate 2 has *rpoB* in addition to *rpoC*. Replicate 1 has mutant variants of all three genes *rpoC, rpoB* and, *rpoA*. *rpoA* has a point mutation A to G (K271E) that has been previously characterized and shown to be crucial for RpoA α-CTD interaction with CRP at some of the CRP dependent promoters [28,29]. These mutants rose in frequency for a period and were in decline by the end of the evolution experiment(Fig. 2).

**Fig 5:**
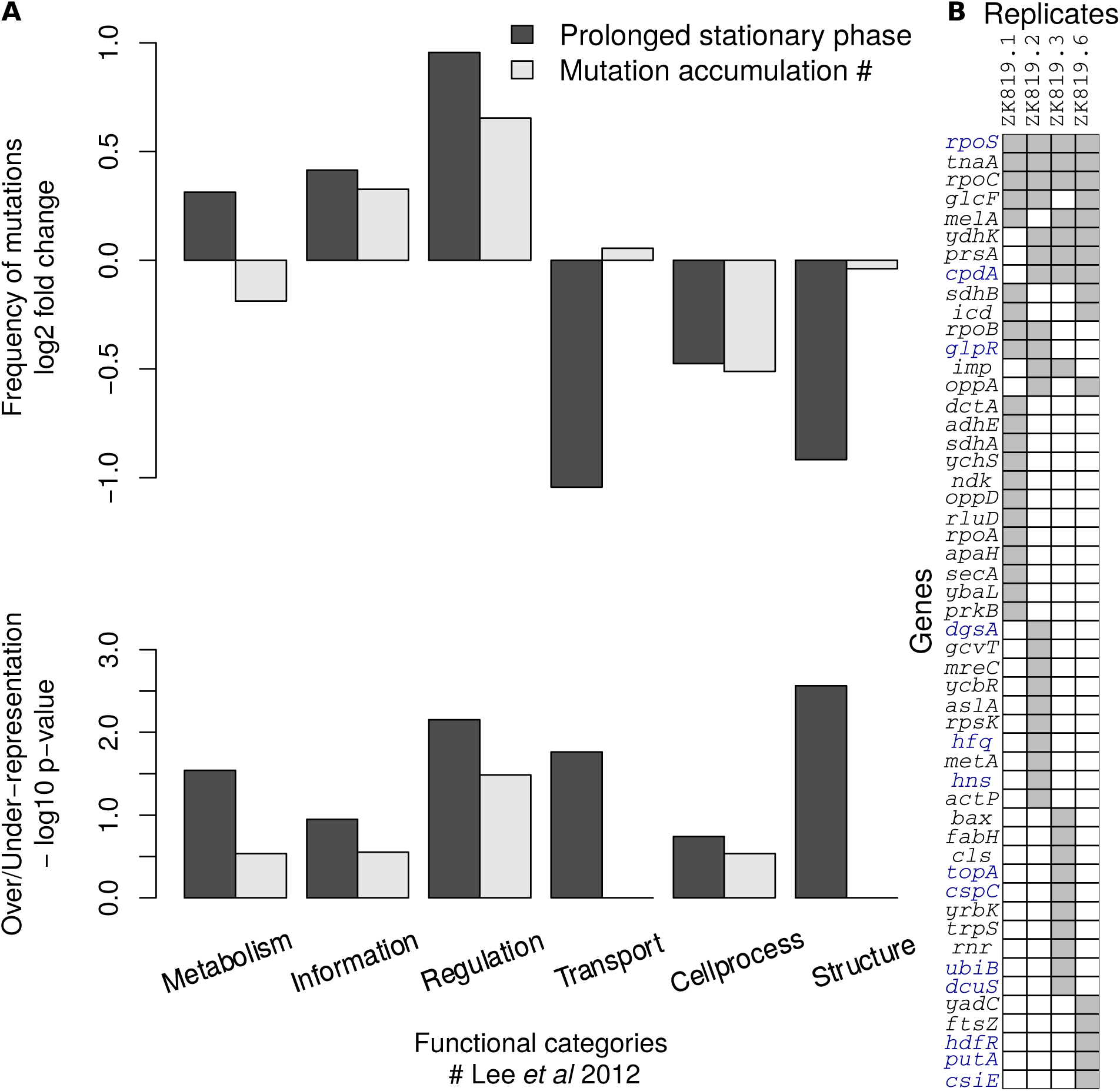
Functional Bias among mutations observed in prolonged stationary phase. (A) the Top plot shows deviation (log2 fold change) from the number of mutations expected in each functional category based on its relative size (total nucleotides) in the genome. The bottom plot shows corresponding -log10 p-values (adjusted for multiple testing with Holm-Bonferroni method) for over/under-representation. Regulation category appears to be over-represented in prolonged stationary phase, no such significant bias was observed for spontaneous mutations under neutral mutation accumulation experiment of Lee *et al* 2012 (B) Binary matrix for genes mutated in four replicates.

Another case of parallel evolution is mutations in the *cpdA* gene. CpdA is a cAMP phosphodiesterase that hydrolysis cAMP [30]. We identified multiple alleles of *cpdA* in four out of five replicates except ZK819.1. We successfully isolated two alleles of *cpdA* from ZK819.3, a T to C point mutation that changes leucine to proline in an alpha helix; and a 3 nucleotides deletion (488-490) that knocks off a histidine residue which is one residue away from critical metal binding histidine residue. Both these *cpdA* alleles were present in the *rpoS819_2* background. These two *cpdA* alleles increase in frequency over time. The 3 nucleotides deletion (488-490) independently appeared in four replicates. These two mutations are likely to negatively affect the function of CpdA.

Another locus of interest is *lptD* (*imp*) gene. LptD is an essential outer membrane protein which along with LptE functions in the assembly of lipopolysaccharides at the surface of outer membrane [31]. A 21 nucleotide deletion at the N-terminal region of *lptD* gene independently appeared in ZK819.2 and ZK819.3. The deletion of 21 nucleotide results in the deletion of seven amino acids. Like *cpdA* alleles, *lptD* alleles appeared during the later stages of the experiment.

## Discussion

Long term stationary phase is a dynamic environment characterized by a multitude of stresses with nutrient limitation as a predominant stress. Previous studies of bacterial evolution in long-term stationary phase have shown the continuous emergence of variants which out-compete their parents [3,8]. A few such variants have been genetically identified [8,9,11,14]. However, much of the variability has been described phenotypically, for example by examination of colony morphotypes [15]. These have indicated that *E. coli* populations in long-term stationary phase are neither homogeneous nor static. In the present study, we have used population genome sequencing to reinforce this at a genetic level.

We show that genetic diversity increases over time in long-term stationary phase populations. It has been suggested that stress could accelerate the generation of genetic diversity by inducing the expression of error-prone DNA polymerases and a reduction in certain repair activities, or by selecting for genetic variants that result in mutator phenotypes [32–42]. In our study, one out of the five populations displayed a strong mutator phenotype. The contribution, if any, of stress-induced mutagenesis to the patterns we observe is an open question. Given the evidence that the expression of various error-prone DNA polymerases is dependent on RpoS [34,36], the fact that our long-term stationary phase cells are deficient in RpoS activity might dampen the argument in favour of selective induction of these DNA polymerases.

Can the level of genetic diversity observed in our study be explained by neutral drift or was it achieved under selection? The role of neutral evolution versus natural selection in the evolution of genetic diversity is a subject of great debate, especially in view of arguments from Michael Lynch that even complex regulatory network architectures can evolve purely by drift [43]. A recent study of mutations in a exponential phase laboratory evolution had used parameters such as the number of non-synonymous vs. synonymous variations to indicate that many of the mutations observed in the study are under selection [44]. The relatively smaller numbers of mutations in our non-mutator lines, and the variability in such a parameter across lines, did not permit a similar analysis. A comparison of the base pair spectrum of mutations in our lines with an experimental model of neutral drift did not reveal any significant signatures of deviation from neutrality. However, a comparison of the genetic diversity that we had observed against what would be predicted by two different models of evolution under pure neutral drift reinforced the view that evolution of genetic diversity in long-term stationary phase is driven by selection and not neutral drift. Further, many non-synonymous mutations we see will result in a switch in the physiochemical characteristics of residues that are conserved to a high degree across bacteria. The prevalence of the repeated occurrence of a large fraction of mutations in certain genes across multiple lines might also be evidence for selection favoring these variants.

We observed a strong enrichment of mutations in regulatory genes. Moreover several of these regulators were independently mutated in more than one replicate populations. This includes further duplications within the *rpoS* gene. One or more RNA polymerase subunit genes are frequently mutated in four of the five replicates suggesting global changes in transcription. A strong case of this genetic parallelism is reflected at the *cpdA* gene. Appearance of various *cpdA* alleles in four of the five populations suggests that there is a strong selection for *cpdA* genetic variants. cAMP-CRP and RpoS controls two largest regulons in *E. coli* by controlling alternate carbon utilization and general stress response during stationary-phase [45–48] However, the interaction between these regulators is unclear. Frequent mutations in *rpoS* and *cpdA* locus in our study may suggest a key regulatory interaction in adaptation under a complex natural environment akin to prolonged stationary-phase. Experiments to evaluate the fitness effect of *rpoS* and *cpdA* mutations and the underlying molecular mechanisms are underway in our laboratory. Evolution by variation in regulatory networks is rapidly gaining ground through various examples in natural populations as well as in laboratory evolution, and the present study provides yet another example to this field.

## Accession Numbers

Genomic deep-sequencing data can be accessed from the NCBI Sequence Read Archive (accession number:SRP094816).

## Acknowledgements

This work was supported by DBT grant:BT/PR3695/BRB/10/979/2011 and Ramanujan fellowship:SR/S2/RJN-49/2010 to A.S.N.S. S.C. is supported by DST-SERB young scientist grant: SB/YS/LS-148/2014. Illumina sequencing was performed at the Centre for Cellular and Molecular Platforms (C-CAMP), Bangalore, India.

